# MPI8 is Potent Against SARS-CoV-2 by Inhibiting Dually and Selectively the SARS-CoV-2 Main Protease and the Host Cathepsin L

**DOI:** 10.1101/2021.06.10.447950

**Authors:** Xinyu R. Ma, Yugendar R. Alugubelli, Yuying Ma, Erol C. Vantasever, Danielle A. Scott, Yuchen Qiao, Ge Yu, Shiqing Xu, Wenshe Ray Liu

**Affiliations:** Texas A&M Drug Discovery Laboratory, Department of Chemistry, Texas A&M University, College Station, TX 77843, USA; Institute of Biosciences and Technology and Department of Translational Medical Sciences, College of Medicine, Texas A&M University, Houston, TX 77030, USA; Department of Biochemistry and Biophysics, Texas A&M University, College Station, TX 77843, USA; Department of Molecular and Cellular Medicine, College of Medicine, Texas A&M University, College Station, TX 77843, USA

## Abstract

A number of inhibitors have been developed for the SARS-CoV-2 main protease (M^Pro^) as potential COVID-19 medications but little is known about their selectivity. Using enzymatic assays, we characterized inhibition of TMPRSS2, furin, and cathepsins B/K/L by more than a dozen of previously developed M^Pro^ inhibitors including MPI1-9, GC376, 11a, 10-1, 10-2, and 10- 3. MPI1-9, GC376 and 11a all contain an aldehyde for the formation of a reversible covalent hemiacetal adduct with the M^Pro^ active site cysteine and 10-1, 10-2 and 10-3 contain a labile ester to exchange with the M^Pro^ active site cysteine for the formation of a thioester. Our data revealed that all these inhibitors are inert toward TMPRSS2 and furin. Diaryl esters also showed low inhibition of cathepsins. However, all aldehyde inhibitors displayed high potency in inhibiting three cathepsins. Their determined IC_50_ values vary from 4.1 to 380 nM for cathepsin B, 0.079 to 2.3 nM for cathepsin L, and 0.35 to 180 nM for cathepsin K. All aldehyde inhibitors showed similar inhibition levels toward cathepsin L. A cellular analysis indicated high potency of MPI5 and MPI8 in inhibiting lysosomal activity, which is probably attributed to their inhibition of cathepsins. Among all aldehyde inhibitors, MPI8 shows the best selectivity toward cathepsin L. With respect to cathepsins B and K, the selective indices are 192 and 150, respectively. MPI8 is the most potent compound among all aldehyde inhibitors in cellular M^Pro^ inhibition potency and anti-SARS-CoV-2 activity in Vero E6 cells. Cathepsin L has been demonstrated to play a critical role in the SARS-CoV-2 cell entry. By selectively inhibiting both SARS-CoV-2 M^Pro^ and the host cathepsin L, MPI8 potentiates dual inhibition effects to synergize its overall antiviral potency and efficacy. Due to its high selectivity toward cathepsin L that reduces potential toxicity toward host cells and high cellular and antiviral potency, we urge serious consideration of MPI8 for preclinical and clinical investigations for treating COVID-19.

## INTRODUCTION

The COVID-19 pandemic has devastated more than 200 countries and territories. As of May 31^st^, 2021, over 170 million people have been confirmed with COVID-19 and the overall death toll has exceeded 3.5 million. Since the identification of the pathogenic virus, SARS-CoV-2, intense efforts have been put into the research of the virus and the development of vaccines and antivirals. There have been quite a few vaccines that showed good effectiveness being approved and distributed. Nevertheless, more and more reported virus variants with mutations on the viral membrane protein Spike could diminish the effectiveness of vaccines and expose humans again to threat in the future. In such a sense, the development of antivirals against SARS-CoV-2 is as important as that of vaccines. Although there are still a lot to learn about SARS-CoV-2 biology and COVID-19 pathogenesis, previous studies of a close related coronavirus SARS-CoV have demonstrated that the main protease (Mpro) is essential to the viral replication and pathogenesis.^1, 2^ Therefore, Mpro is considered as a valid antiviral target for SARS-CoV-2. Unlike Spike that is highly mutable, Mpro is highly conserved as exemplified by the 96% sequence identity shared between its genes from SARS-CoV and SARS-CoV-2.^3^ Small molecules that are developed for Mpro from SARS-CoV-2 might serve as general antivirals for coronaviruses. So far, a number of Mpro inhibitors have been developed.^4-11^ Some have shown promising *in vitro* and *in vivo* efficacy in inhibiting SARS-CoV-2.^8^

Mpro is a cysteine protease. Its enzymatic activity relies on the catalytic Cys145 residue at the active site. Many currently developed Mpro inhibitors contain an aldehyde or α-ketoamide warhead that reacts reversibly with Cys145 to generate a covalent hemiacetal intermediate. A lot of these inhibitors are also peptidomimetics that were derived from Mpro substrates. The nonselective reactivity of a keto group toward a thiol or hydroxyl group and the peptidomimetic structure make many Mpro inhibitors susceptible to interact with cysteine and serine proteases in host cells as well. Although this propensity leaves a selectivity concern for these Mpro inhibitors, it could also create the potential for the inhibition of host proteases that serve critical roles in the SARS-CoV-2 replication and pathogenesis. Accumulative evidence has shown that certain host proteases prime the SARS-CoV-2 Spike protein for viral packaging, interactions with ACE2, and viral entry into the host. These include two serine proteases furin and transmembrane protease serine 2 (TMPRSS2) and a cysteine protease cathepsin L.^12-15^ Small molecule medications that inhibit furin, TMPRSS2 and cathepsin L have shown efficacy in inhibiting SARS-CoV-2.^12-14^ Some of them including camostat and K777 are under clinical trials for COVID-19.^16, 17^ An Mpro inhibitor that dually inhibits a host protease for the virus replication and pathogenesis potentiates improved overall antiviral potency and efficacy. To explore potential dual inhibition of a critical host protease, we tested inhibition potency of fourteen previously reported Mpro inhibitors including eleven peptidomimetic aldehydes and three diaryl esters against furin, TMPRSS2 and cathepsin L. To explore selectivity of these Mpro inhibitors against other host cysteine proteases, we tested their inhibition of cathepsins B and K as well. Our results showed that a previously developed Mpro inhibitor MPI8 dually inhibits Mpro and cathepsin L with high potency and selectivity.

## RESULTS

In our previous work, we reported MPI1-9 shown in Figure 1 as a series of potent peptidyl aldehyde inhibitors for Mpro, with the most potent one having a single-digit nanomolar IC_50_ value.^5^ In the following cytopathic effect (CPE) assays, two compounds, MPI5 and MPI8 showed low micromolar EC_100_ values against SARS-CoV-2 in Vero E6 cells and submicromolar EC_100_ values against SARS-CoV-2 in ACE2^+^ A549 cells, while other MPIs displayed much lower antiviral potencies despite that they have much lower enzymatic inhibition IC_50_ values compared to MPI5 and MPI8. Vero E6 cells were derived from an African monkey. A549 cells are adenocarcinomic human alveolar basal epithelial cells. The higher antiviral efficacy of MPI5 and MPI8 in ACE2^+^ A549 cells compared to that in Vero E6 cells implies that these inhibitors might interfere with a human pathway critical for viral replication in addition to Mpro inhibition. As peptidyl aldehydes, MPIs are very likely to covalently inhibit human cysteine or serine proteases. Since furin, TMPRSS2 and cathepsin L are known to serve functions in the SARS-CoV-2 replication, it is possible that the observed potent antiviral efficacies of MPI5 and MPI8 in ACE^+^ A549 cells were a result of dual-inhibition of Mpro and a host protease associated with SARS-CoV-2 replication. To test this potential, we characterized MPI1-9 on their inhibition of furin, TMPRSS2 and cathepsin L. We characterized their inhibition of cathepsins B and K as well for selectivity comparison.

**Figure 1.**
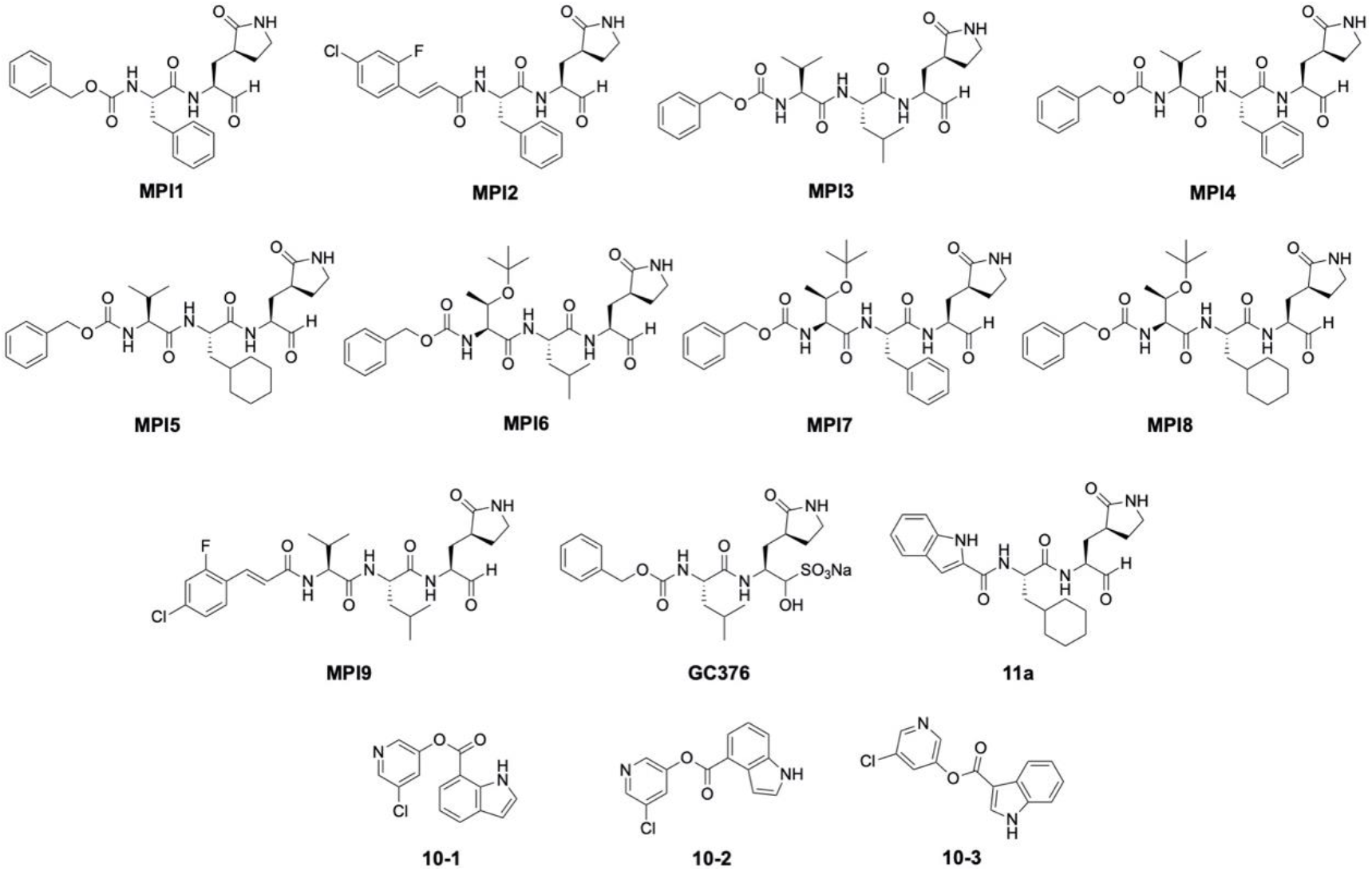
Structures of the tested compounds in this study, MPI1-9, 11a, GC376, 10-1, 10-2, and 10-3.

we first determined enzymatic inhibition IC_50_ values of MPI1-9 against furin and TMPRSS2 using kinetic assays where the rate of the hydrolysis of a fluorogenic substrate by furin or TMPRSS2 was measured. The protocols were adapted and modified from published literatures.^18- 20^ Both substrates and enzymes were commercially acquired. The substrates that we used for the inhibition assay of fursin and TMPRSS2 were Pyr-Arg-Thr-Lys-Arg-7-amino-4-methylcoumarin (AMC) for furin and Boc-Gln-Ala-Arg-AMC for TMPRSS2. Both substrates were titrated to a final concentration of 10 µM in the assay. In the assay, enzymes were titrated to a final concentration of 10 nM for furin and 1 µM for TMPRSS2 so that an optimal signal -to-noise ratio could be reached. MPI1-9 potentially react with the active site serine of both furin and TMPRSS2 to form a hemiacetal (Figure 2A). We first screened MPI1-9 at a final concentration of 1 μM to see whether they inhibit furin and TMPRSS2. All experiments were conducted in triplicates. As shown in Figures 2B and 2C, all nine molecules did not exhibit significant inhibition of both enzymes at 1 μM. In a separate study, we developed a cellular Mpro inhibition assay for analyzing cellular Mpro inhibition potencies of inhibitors.^21^ This assay revealed that MPI8 inhibited Mpro in a human host cell with an IC_50_ value of 31 nM. This cellular potency was corroborated by a determined antiviral EC_50_ value of 30 nM for MPI8 in Vero E6 cells. Due to this potent cellular and antiviral activities at a nanomolar level for MPI8, we deemed that measuring related compounds in their micromolar concentrations for inhibition of both furin and TMPRSS2 is not meaningful since a high micromolar concentration of these compounds will not be used in real antiviral tests. Based on our results with respect to furin and TMPRSS2, we conclude that MPI1-9 do not inhibit both enzymes.

**Figure 2.**
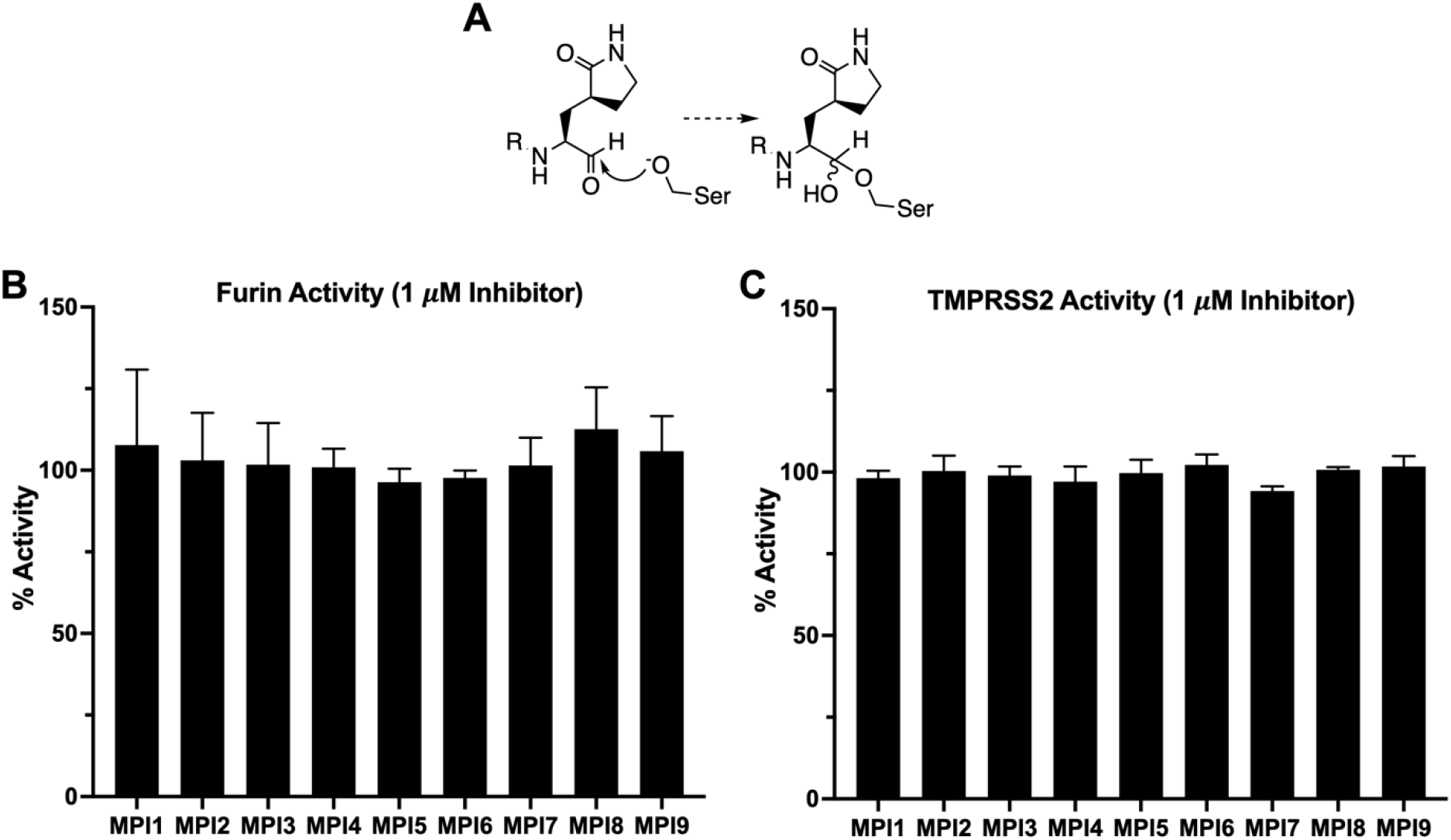
(A) A scheme shown the reversible covalent interaction between aldehyde inhibitors with serine proteases. (B) Percentage of Furin activity after treatment of MPI1-9 at 1 µM concertation. (C) Percentage of TMPRSS2 activity after treatment of MPI1-9 at 1 µM concertation.

To analyze inhibition potencies of MPI1-9 toward cathepsins B, K, and L, we adapted a literature protocol and used a fluorogenic substrate Z-Phe-Arg-AMC.^22^ This substrate was titrated to a final concentration of 20 μM in the final assay. For the three enzymes, their final concentrations used in the inhibition assay were 2 nM for cathepsin L, 1 nM for cathepsin K, and 5 nM for cathepsin B. As shown in Figure 3 and Table 1, MPI1-9 potently inhibit cathepsin L. Their determined IC_50_ values range from 79 pm for MPI1 and 2.3 nm for MPI5. MPI5 and MPI8 are two weakest cathepsin L inhibitors among all MPIs. We noticed that the determined IC_50_ value for MPI1 was significantly lower than the concentration of cathepsin L used in the IC_50_ characterization assay. We acquired cathepsin L from a commercial source. Cathepsin L is known to be processed from a proprotein. We think that it is highly probable that the active cathepsin L component was significantly less than the overall enzyme concentration. In comparison to other MPIs, MPI1 is much smaller and less rigid than MPI2. Its small size and structural flexibility may contribute to its potent inhibition of cathepsin L. In comparison to inhibiting cathepsin L, MPI1-9 displayed much higher IC_50_ values in inhibiting cathepsin B, ranging from 4.1 nM for MPI2 to 380 nM for MPI6. From the results, all MPIs are selective toward cathepsin L in comparison to cathepsin B. The selective indices vary from 22 for MPI2 and 840 for MPI6. As the molecule with the highest cellular and antiviral potencies among all MPIs, MPI8 has a selective index of 192 in targeting cathepsin L over cathepsin B. In comparison to inhibiting cathepsin L, MPI1-9 displayed also higher IC_50_ values in inhibiting cathepsin K, ranging from 0.78 nM for MPI2 and 180 nM for MPI8. All MPIs are selective toward cathepsin L in comparison to cathepsin K. The selective indices vary from 2.4 for MPI6 to 150 for MPI8. By judging selectivity toward cathepsin L over both cathepsin B and cathepsin K, MPI8 has the best characteristics. Other MPIs that displayed modest to strong selectivity toward cathepsin L over both cathepsin B and cathepsin L are MPI4, MPI5 and MPI7.

**Table 1.**
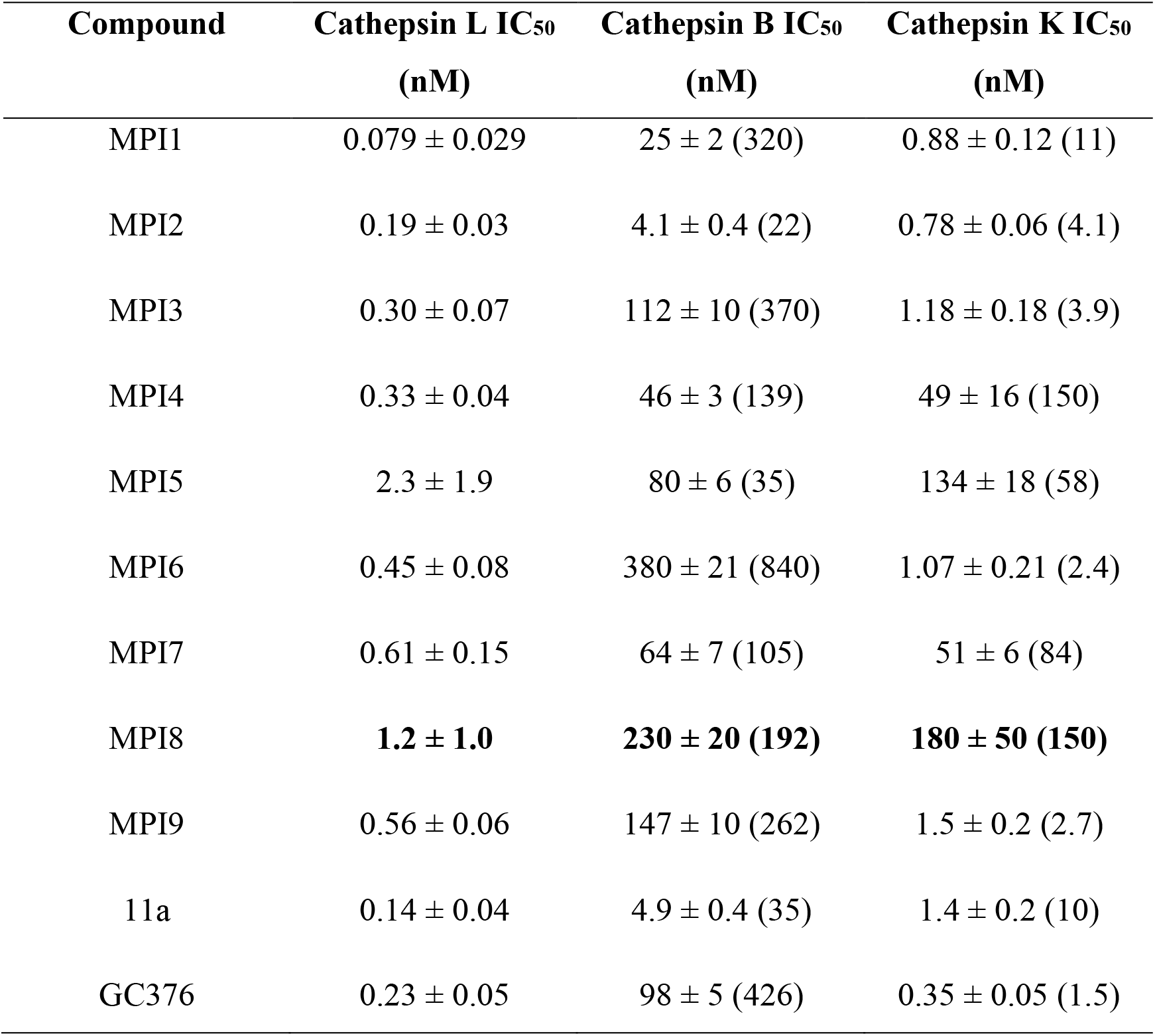
IC_50_ of MPI1-9, 11a, and GC376 against cathepsin L, cathepsin B, and cathepsin K. Parenthesized numbers indicate selective indices with respect to the activity toward cathepsin L. values for MPI8 are highlighted.

**Figure 3.**
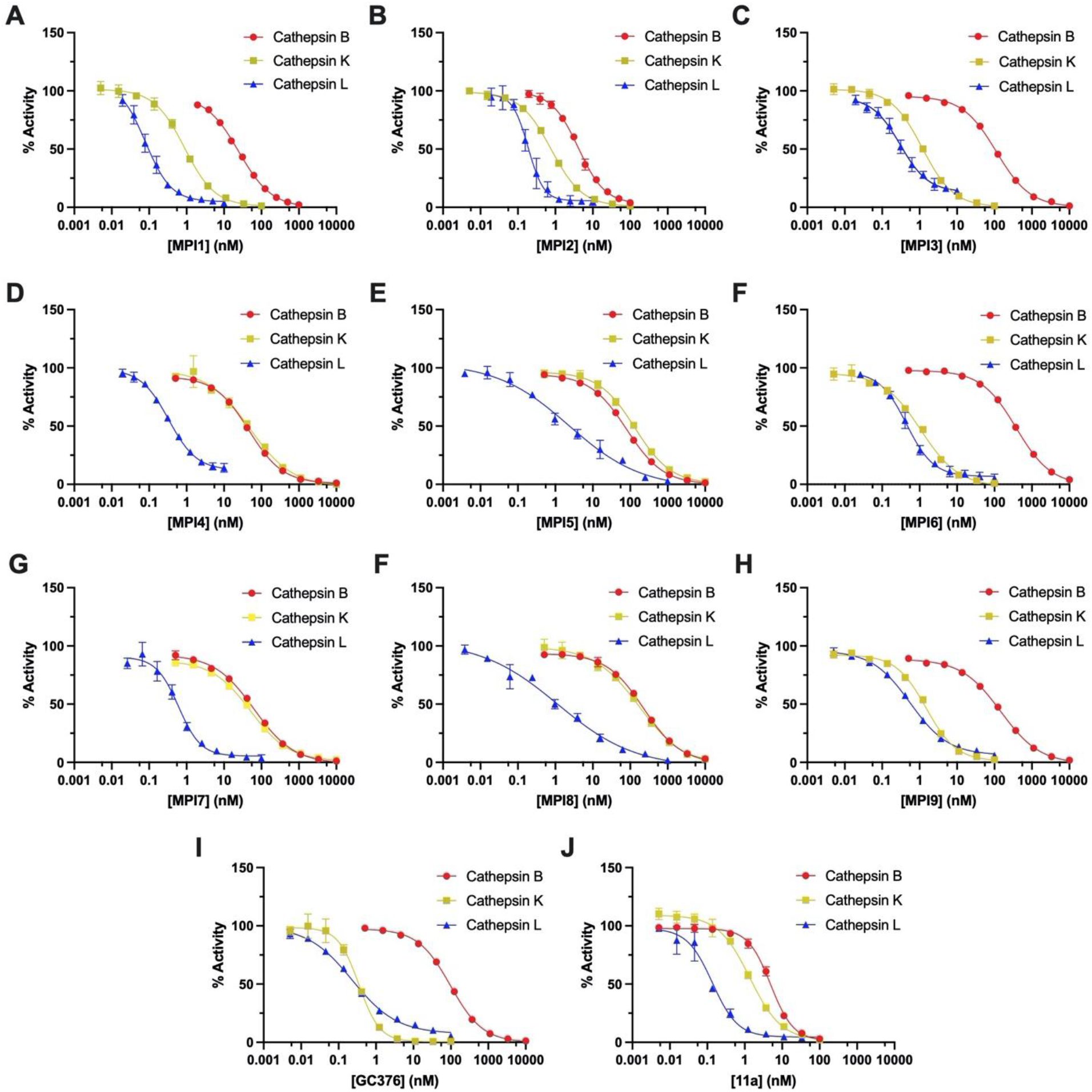
IC_50_ curves for MPI1-9, GC376, and 11a against cathepsin L, cathepsin B or cathepsin K.

GC376^23^ and 11a^7^ are two investigational drugs undergoing clinical trials for the treatment of COVID-19 patients in United States. They are also peptidyl aldehydes and share structural similarity to our MPI1-9 molecules. We acquired GC376, synthesized 11a and then conducted their IC_50_ determination for furin, TMPRSS2, and cathepsins B, K and L. Both GC376 and 11a showed no inhibition of furin and TMPRSS2 at 1 μM. However, both molecules displayed high potencies in inhibiting all three cathepsins (Figure 3 and Table 1). Both 11a and GC376 are selective toward cathepsin L over cathepsins B and K. However, selective indices for 11a are modest. Its selectivity index toward cathepsin L over cathepsin K is only 10. GC376 has very poor selectivity toward cathepsin L over cathepsin K with a selectivity index of only 1.5.

In a previous study, we synthesized three diaryl esters 10-1, 10-2 and 10-3 and determined their IC_50_ values in inhibiting Mpro as 0.067, 0.038, and 7.6 μM, respectively. All three molecules exhibited also cellular potencies in inhibiting Mpro. We measured these three compounds in their inhibition of furin, TMPRSS2 and cathepsins B, K and L as well. All three compounds showed no inhibition of furin and TMPRSS2. They did not display significant inhibition of cathepsins B, K and L up to the concentration of 10 μM (Figure 4).

**Figure 4.**
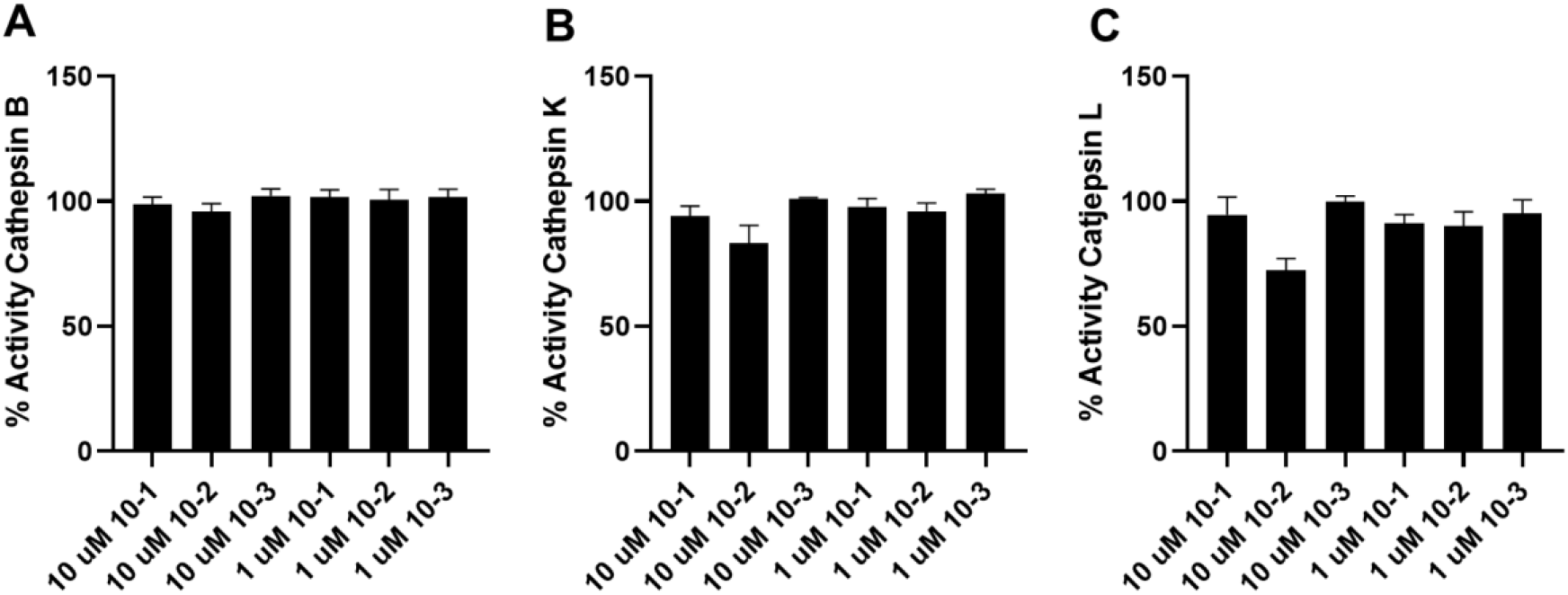
Indole chloropyridinyl esters showed little inhibition against cathepsin L, cathepsin B or cathepsin K. (A) Percentage of cathepsin B activity after treatment with 10-1, 10-2, and 10-3. (B) Percentage of cathepsin K activity after treatment with 10-1, 10-2, and 10-3. (C) Percentage of cathepsin L activity after treatment with 10-1, 10-2, and 10-3.

**Figure 5.**
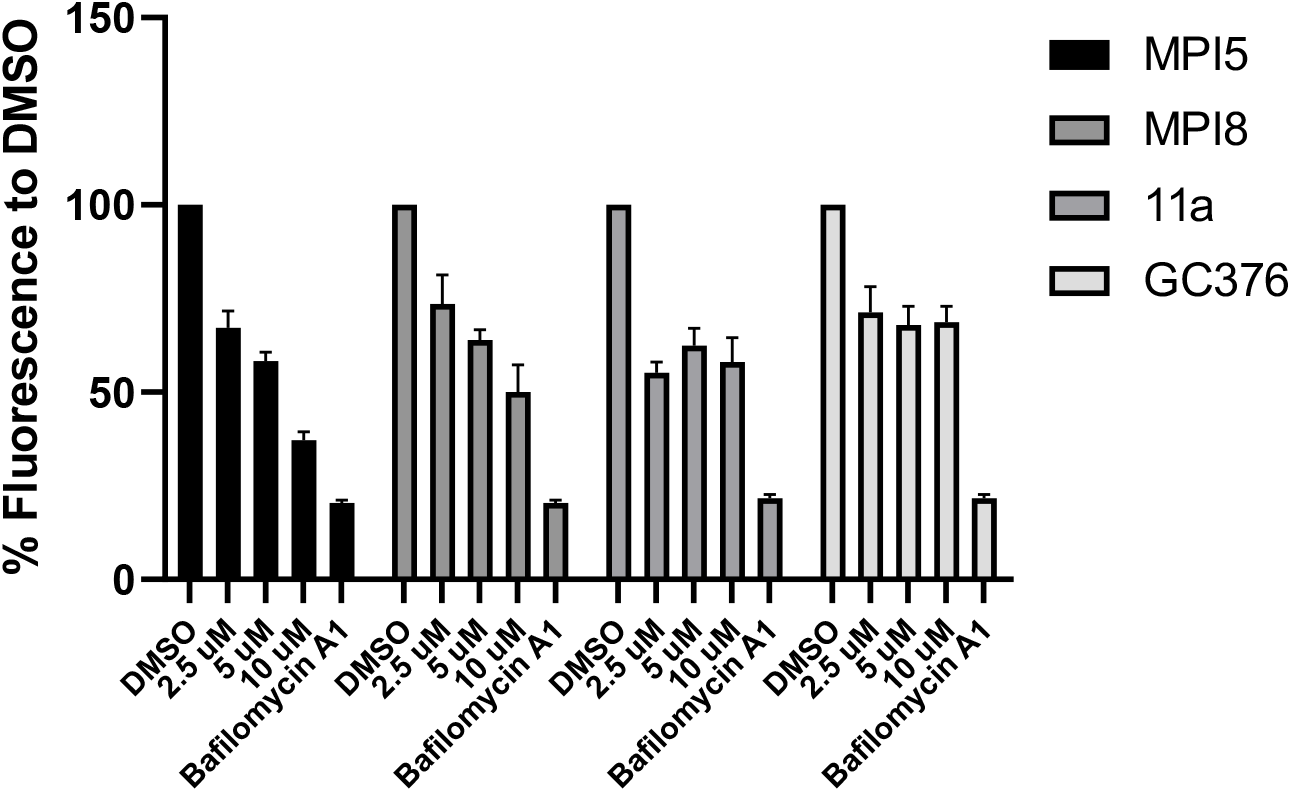
Inhibition of cellular lysosomal activity by MPI5, MPI8, 11a, and GC376. The intensity of cellular fluorescence indicated how much a fluorogenic substrate was degraded by lysosomes, representing cellular lysosomal activity. For each compound, three concentrations (10 µM, 5 µM, and 2.5 µM) were tested and the fluorescence signals were normalized to that of DMSO group.

Among all MPIs, MPI5 and MPI8 showed highest potencies in inhibiting SARS-CoV-2 in Vero E6 cells with EC_50_ values of 73 and 30 nM, respectively^21^. MPI8 has the best selectivity indices toward cathepsin L over cathepsins B and K among all compounds we tested. Although not comparable to MPI8, MPI5 displays modest cathepsin L selectivity indices over cathepsins and K. Due to the known role of cathepsin L in the SARS-CoV-2 replication and pathogenesis, both MPI5 and MPI8 likely exert dual functions in inhibiting both Mpro and cathepsin L to convene strong antiviral potencies against SARS-CoV-2. The modest to high cathepsin L selectivity indices over cathepsins B and K will also make the two MPIs less toxic to host cells. However, MPI5 and MPI8 are two least potent cathepsin L inhibitors among all aldehyde compounds we tested, leaving it questionable whether MPI5 and MPI8 inhibit genuinely cathepsin L activity, or more generally, lysosomal or endosomal activities in cells as they do *in vitro*. To confirm that, we further used HEK293T cells to perform an intracellular lysosomal activity assay in the presence of MPI5, MPI8, 11a, or GC376. All four compounds showed SARS-CoV-2 inhibition efficacy at the cellular level. Cells were first treated with 10, 5 or 2.5 μM of an inhibitor. Then a commercially available self-quenched fluorogenic substrate (ab234622 from Abcam) was provided to assess the lysosomal activity of treated cells. Lysosomotropic agent bafilomycin A1, which inhibits lysosomal activity, was used as control. The results showed that MPI5, MPI8, 11a, and GC376 all inhibited cellular lysosomal activity at a concentration as low as 2.5 µM (Figure 3), strongly suggesting that all these compounds inhibit SARS-CoV-2 replication in cells via a dual-target mechanism.

## DISCUSSION

It is quite intriguing that all peptidyl aldehyde inhibitors we tested showed remarkable potency in the inhibition of cathepsins B, K and L *in vitro*, especially cathepsin L. This observation might not be pure coincidence but have underlying connections with the substrate specificity of cathepsin L and Mpro. According to Mpro crystal structure,^24^ glutamine residue or γ-lactam residue are considered the best fit for its S1 pocket, which overlaps with the substrate specificity of cathepsin L in which glutamine and glycine are the most favored residues at P1 position.^25^ At P2 position, Mpro favors leucine the most and accommodates many other hydrophobic residues as well.^26^ This also overlaps with P2 specificity of cathepsin L of aliphatic residues. The overlapped substrate specificity between these two proteases makes peptidomimetic inhibitors designed based on the substrate specificity of one of them likely to inhibit the other one as well. This implies also that a non-peptidomimetic inhibitor should be more selective between Mpro and cathepsin L. As we demonstrated for 10-1, 10-2 and 10-3, all three diaryl esters potently inhibit Mpro but showed close to no inhibition of cathepsins B, K and L up to 10 μM. For the development of selective Mpro inhibitors, diaryl esters is a direction worth pursuing. In the current study, we demonstrated also that MPI5, MPI8, 11a, and GC376 inhibit intracellular lysosomal activity, unveiling more complicated pharmacology of these peptidyl aldehyde compounds in cells. From one perspective, by inhibiting the activity of cathepsin L and lysosomes in cells, these Mpro inhibitors can also block SARS-CoV-2 entry, providing them with a dual mechanism-of-action other than mono-specific inhibition of Mpro activity. One obvious advantage of the dual-target mechanism is the less susceptibility to mutation of virus and consequent drug resistance. However, from the other perspective, the inclusion of a human protease as an immediate drug target also means a higher chance of adversary effects and may significantly lower their therapeutic indexes, limiting their potential as an anti-SARS-CoV-2 therapy. As far as we know, cathepsin L is the only enzyme in the cathepsin family that showed clear relevance to SARS-CoV-2 entry. This makes it particularly a concern for those inhibitors that showed relatively poor selectivity between different cathepsins. Among the four compounds that showed anti-lysosomal activity in cells, GC376 has barely any selectivity between cathepsin K and cathepsin L, 11a and MPI5 have some moderate selectivity, and MPI8 is the most selective one despite its relatively low potency toward cathepsin L, which implies that MPI8 is a better candidate for the dual-target inhibition of SARS-CoV-2. Among all aldehyde-based Mpro inhibitors, MPI8 has the best cellular Mpro inhibition and antiviral potencies. Its high selectivity toward the inhibition of cathepsin L in host cells provides dual functions to inhibit SARS-CoV-2 with less off-target concerns than other aldehyde-based inhibitors. Given all evidences accumulated so far, we cautiously urge preclinical and clinical investigation of MPI8 in the treatment of COVID-19.

### Conclusion

In this study, we characterized inhibition of furin, TMPRSS2 and cathepsins B, K and L by 11 peptidyl aldehyde and 3 diaryl ester Mpro inhibitors. All peptidyl aldehydes potently inhibit cathepsins especially for cathepsin L that is a known host protease playing a key role in the SARS-COV-2 replication and pathogenesis. Four peptidyl aldehydes MPI5, MPI8, 11a, and GC376 with antiviral potencies showed also inhibition of lysosomal activity in HEK293T cells, implying that these inhibitors have a dual-target mechanism of action. Among them, MPI8 has the most cellular Mpro inhibition and antiviral potency and a high selectivity toward inhibiting cathepsin L over cathepsins B and K, making it the best candidate to exert dual functions to inhibit SARS-CoV-2. In contrast, diaryl esters 10-1, 10-2, and 10-3 showed excellent selectivity between Mpro and the human proteases, indicating that the development of diaryl esters could be a general route to develop selective Mpro inhibitors. Overall, given its high cellular Mpro inhibition and anti-SARS-CoV-2 potency as well as high selectivity toward cathepsin L over other human proteases, we urge serious consideration of MPI8 for preclinical and clinical investigations for treating COVID-19.

## Methods

### Inhibition assay for cathepsin L

The assay was performed in the following assay buffer: 100 mM MES-NaOH solution (pH 5.5) containing 2.5 mM EDTA, 2.5 mM DTT and 9% DMSO. The stock solution of the enzyme was diluted to 20 nM with assay buffer. Stock solutions of inhibitors were prepared in DMSO. A 10 mM stock solution of the fluorogenic substrate Z-Phe-Arg-AMC was diluted with assay buffer. The final concentration in enzymatic assay of DMSO was 10%, and those of the substrate and cathepsin L were 20 µM and 2 nM, respectively. Into a well containing 39 µL assay buffer, 1 µL inhibitor solution (or DMSO) and 10 µL diluted solution of cathepsin L were added and mixed thoroughly, and then incubated at 37 °C for 30 min. The reaction was initiated by adding 50 µL diluted solution of substrate and the fluorescence intensity at 440 nm under 360 nm excitation was measured. Experiments were performed in triplicate with at least ten different concentrations of each inhibitor and both positive and negative controls. The initial rate was calculated according to the fluorescent intensity in the first five minutes by linear regression, which was then normalized according to the initial rate of positive and negative controls. IC_50_ curve was simulated by GraphPad 8.0 using sigmoidal model (four parameters).

### Inhibition assay for cathepsin B

The assay was performed in the following assay buffer: 100 mM MES-NaOH solution (pH 6.0) containing 2.5 mM EDTA, 2.5 mM DTT, 0.001% Tween 20 and 9% DMSO. The stock solution of the enzyme was diluted to 50 nM with assay buffer. Stock solutions of inhibitors were prepared in DMSO. A 10 mM stock solution of the fluorogenic substrate Z-Phe-Arg-AMC was diluted with assay buffer. The final concentration in enzymatic assay of DMSO were 10%, and those of the substrate and cathepsin B was 20 µM and 5 nM, respectively. Into a well containing 39 µL assay buffer, 1 µL inhibitor solution (or DMSO) and 10 µL diluted solution of cathepsin B were added and mixed thoroughly, and then incubated at 37 °C for 30 min. The reaction was initiated by adding 50 µL diluted solution of substrate and the fluorescence intensity at 440 nm under 360 nm excitation was measured. Experiments were performed in triplicate with at least ten different concentrations of each inhibitor and both positive and negative controls. The initial rate was calculated according to the fluorescent intensity in the first five minutes by linear regression, which was then normalized according to the initial rate of positive and negative controls. IC_50_ curve was simulated by GraphPad 8.0 using sigmoidal model (four parameters).

### Inhibition assay for cathepsin K

The assay was performed in the following assay buffer: 100 mM MES-NaOH solution (pH 5.5) containing 2.5 mM EDTA, 2.5 mM DTT and 9% DMSO. The stock solution of the enzyme was diluted to 10 nM with assay buffer. Stock solutions of inhibitors were prepared in DMSO. A 10 mM stock solution of the fluorogenic substrate Z-Phe-Arg-AMC was diluted with assay buffer. The final concentration in enzymatic assay of DMSO was 10%, and those of the substrate and cathepsin K were 20 µM and 1 nM, respectively. Into a well containing 39 µL assay buffer, 1 µL inhibitor solution (or DMSO) and 10 µL diluted solution of cathepsin K were added and mixed thoroughly, and then incubated at 37 °C for 30 min. The reaction was initiated by adding 50 µL diluted solution of substrate and the fluorescence intensity at 440 nm under 360 nm excitation was measured. Experiments were performed in triplicate with at least ten different concentrations of each inhibitor and both positive and negative controls. The initial rate was calculated according to the fluorescent intensity in the first five minutes by linear regression, which was then normalized according to the initial rate of positive and negative controls. IC_50_ curve was simulated by GraphPad 8.0 using sigmoidal model (four parameters).

### Inhibition assay for furin

The assay was performed in the following assay buffer: 100 mM HEPES buffer (pH 7.0) containing 0.2% Triton X-100, 2 mM CaCl_2_, 0.02% sodium azide, and 1 mg/mL BSA. The stock solution of the enzyme was diluted to 100 nM with assay buffer. Stock solutions of inhibitors were prepared in DMSO. A 10 mM stock solution of the fluorogenic substrate Pyr-Arg-Thr-Lys-Arg-AMC was diluted with assay buffer. The final concentrations in enzymatic assay of the substrate and furin were 10 µM and 10 nM, respectively. Into a well containing 39 µL assay buffer, 1 µL inhibitor solution (or DMSO) and 10 µL diluted solution of furin were added and mixed thoroughly, and then incubated at 37 °C for 30 min. The reaction was initiated by adding 50 µL diluted solution of substrate and the fluorescence intensity at 440 nm under 360 nm excitation was measured. Experiments were performed in triplicate with different concentrations of each inhibitor and both positive and negative controls. The initial rate was calculated according to the fluorescent intensity in the first five minutes by linear regression, which was then normalized according to the initial rate of positive and negative controls. IC_50_ curve was simulated by GraphPad 8.0 using sigmoidal model (four parameters).

### Inhibition assay for TMPRSS2

The assay was performed in the following assay buffer: 50 mM Tris (pH 8.0) containing 150 mM NaCl, and 0.01% Tween20. The stock solution of the enzyme was diluted to 10 µM with assay buffer. Stock solutions of inhibitors were prepared in DMSO. A 10 mM stock solution of the fluorogenic substrate Boc-Gln-Ala-Arg-AMC was diluted with assay buffer. The final concentrations in enzymatic assay of the substrate and TMPRSS2 were 10 µM and 1 µM, respectively. Into a well containing 39 µL assay buffer, 1 µL inhibitor solution (or DMSO) and 10 µL diluted solution of TMPRSS2 were added and mixed thoroughly, and then incubated at 37 °C for 30 min. The reaction was initiated by adding 50 µL diluted solution of substrate and the fluorescence intensity at 440 nm under 360 nm excitation was measured. Experiments were performed in triplicate with different concentrations of each inhibitor and both positive and negative controls. The initial rate was calculated according to the fluorescent intensity in the first five minutes by linear regression, which was then normalized according to the initial rate of positive and negative controls. IC_50_ curve was simulated by GraphPad 8.0 using sigmoidal model (four parameters).

### Intracellular lysosomal activity assay

The protocol was adapted from that provided by Abcam with some modifications. HEK293T cells were grown in DMEM media containing 10 % FBS in standard 24-well plate and incubated under 37 °C, 5 % CO_2_ overnight. Media was then removed and replaced with fresh DMEM media containing different concentrations of test compounds with 0.1 % DMSO (experimental group), 0.1 % DMSO (negative control group) or 1 × bafilomycin A1 (positive control group). The cells were incubated under 37 °C, 5 % CO_2_ for 2 hours. The media was then removed and replaced with fresh DMEM media containing 0.5 % FBS and the same concentration of test compounds, DMSO or bafilomycin. 15 µL of self-quenched fluorogenic substrate was added to each well per 1 mL of media. The cells were incubated under 37 °C, 5 % CO_2_ for another 2 hours. The media was then removed, and the cells were harvested, washed with 1 mL ice cold assay buffer (provided in assay kit) twice and resuspended in PBS buffer for flow cytometry analysis under 488 nm excitation. Experiments were performed in triplicate.

